# Tardigrade community microbiomes in North American orchards include putative endosymbionts and plant pathogens

**DOI:** 10.1101/2022.01.28.478239

**Authors:** Laura E. Tibbs-Cortes, Bienvenido W. Tibbs-Cortes, Stephan Schmitz-Esser

## Abstract

The microbiome of tardigrades, a phylum of microscopic animals best known for their ability to survive extreme conditions, is poorly studied worldwide and completely unknown in North America. An improved understanding of tardigrade-associated bacteria is particularly important because tardigrades have been shown to act as vectors of the plant pathogen *Xanthomonas campestris* in the laboratory. However, the potential role of tardigrades as reservoirs and vectors of phytopathogens has not been investigated further. This study analyzed the microbiota of tardigrades from six apple orchards in central Iowa, USA, and is the first analysis of the microbiota of North American tardigrades. It is also the first ever study of the tardigrade microbiome in an agricultural setting. We utilized 16S rRNA gene amplicon sequencing to characterize the tardigrade community microbiome across four contrasts: location, substrate type (moss or lichen), collection year, and tardigrades versus their substrate. Alpha diversity of the tardigrade community microbiome differed significantly by location and year of collection but not by substrate type. Our work also corroborated earlier findings, demonstrating that tardigrades harbor a distinct microbiota from their environment. We also identified tardigrade-associated taxa that belong to genera known to contain phytopathogens (*Pseudomonas, Ralstonia*, and the *Pantoea/Erwinia* complex). Finally, we observed members of the genera *Rickettsia* and *Wolbachia* in the tardigrade microbiome; because these are obligate intracellular genera, we consider these taxa to be putative endosymbionts of tardigrades. These results suggest the presence of putative endosymbionts and phytopathogens in the microbiota of wild tardigrades in North America.

## 1 Introduction

Tardigrades are a poorly-studied but globally ubiquitous phylum of microscopic animals. They are members of the superphylum Ecdysozoa, a group that also includes arthropods and nematodes. All tardigrades are aquatic; however, while some live in bodies of fresh or salt water, they are most commonly collected from moss or lichen, where they live in interstitial films of water. When this water dries up, tardigrades survive by dehydrating and entering a state of dramatically reduced metabolism known as cryptobiosis (Kinchin 1994). In this state, they are famously able to survive extreme conditions, ranging from temperatures near absolute zero (Becquerel 1950) to the vacuum of space (Jönsson et al. 2008). Despite extensive study of tardigrades’ survival abilities, little is known about many aspects of their biology, including their microbiota (Vecchi et al. 2018). This is particularly important because tardigrades’ presence in moss and lichen, which often grow on tree bark, brings them into close contact with trees, including important orchard crops such as apple trees (*Malus domestica* L. Borkh). Therefore, any plant pathogens present in the tardigrade microbiota have the potential to affect these crops, underscoring the importance of understanding the tardigrade microbiota in an agricultural context.

The first study of tardigrade-associated bacteria, published in 1999, found that bacteria of the phytopathogenic genus *Xanthomonas* could be grown from the feces of the tardigrade *Macrobiotus hufelandi* C.A.S. Schultze, 1834 isolated from the wild. However, attempts to inoculate *Mac. hufelandi* with *Serratia marcescens* were unsuccessful, suggesting a non-random relationship between *Mac. hufelandi* and *Xanthomonas* (Krantz, Benoit, and Beasley 1999). The following year, a second paper showed that *Mac. hufelandi* exposed to infected leaves could spread *X. campestris* pv. *raphani* (the causal pathogen of radish leaf spot disease) to healthy radish seedlings in the laboratory. This demonstrated that *Mac. hufelandi* can act as a vector of radish leaf spot disease (Benoit et al. 2000).

Animal vectors are known to spread many plant diseases, with major consequences for crop production worldwide (Ng and Falk 2006; Mew 1993; Duveiller, Bragard, and Marite 1997). Current research focuses on insect vectors (Ng and Falk 2006), but the work of Krantz et al. and Benoit et al. demonstrate that at least one tardigrade species (*Mac. hufelandi*) can spread bacterial disease in plants (Benoit et al. 2000) and can act as reservoirs of plant pathogens (Krantz, Benoit, and Beasley 1999). Because *Mac. hufelandi* and many other tardigrade species live in close contact with plants, bacteria deposited in their feces may infect these plants, especially as other Ecdysozoans are known to spread phytopathogens in this manner (Dutta et al. 2014; Stavrinides, McCloskey, and Ochman 2009). Tardigrades also have the potential to spread phytopathogenic bacteria over large areas because many tardigrade species are cosmopolitan (Meyer 2013) and may be dispersed by wind or migratory birds (Mogle et al. 2018).

The genus *Xanthomonas* found in association with *Mac. hufelandi* includes pathovars that infect staple food crops including rice (*Oryza sativa* L.) (Mew 1993), wheat (*Triticum aestivum* L.) (Duveiller, Bragard, and Marite 1997), and maize (*Zea mays* L.) (Karamura et al. 2007) with potentially devastating effects. For example, bacterial blight (*X. oryzae* pv. *oryzae*) can cause yield losses of up to 50% in rice infected as seedlings, impacting both economies and food security (Mew 1993). Yet, although tardigrades are known vectors of this important genus, there has been no additional literature published on phytopathogens associated with tardigrades in the past two decades.

Recent studies of the tardigrade microbiome, while not focusing on phytopathogens, have leveraged advances in sequencing technology by using 16S rRNA gene amplicon sequencing. Vecchi *et al*. surveyed the microbial communities associated with six tardigrade species: *Acutuncus antarcticus* (Richters, 1904) collected from freshwater sediment in Antarctica, after which a subsample was raised in laboratory culture; *Ramazzottius oberhaeuseri* (Doyère, 1840), collected from lichen on two different trees in Italy; *Macrobiotus macrocalix* Bertolani & Rebecchi, 1993 and *Richtersius coronifer* (Richters, 1903), both collected from the same moss on a rock in Sweden; and *Echiniscus trisetosus* Cuénot, 1932, and *Paramacrobiotus areolatus* (Murray, 1907), both collected from the same moss on a rock in Italy. The authors found that the tardigrade microbiome is dominated by *Proteobacteria* and *Bacteroidetes*, is distinct from and usually less diverse than that of their substrates, differs among tardigrade species, and is altered by laboratory culturing of the tardigrades. Vecchi *et al*. also identified potential endosymbionts of the obligate intracellular order *Rickettsiales* within the tardigrade microbiome (2018). This is particularly intriguing because the genera *Wolbachia* and *Rickettsia*, both members of *Rickettsiales*, are known to have reproductive effects on their hosts, including inducing parthenogenesis (Giorgini et al. 2010; Werren, Baldo, and Clark 2008). Notably, parthenogenesis is common in tardigrades (Bertolani 2001; Guil et al. 2022). A subsequent analysis of these data identified four putative endosymbionts in the order *Rickettsiales*, three of which belonged to *Anaplasmataceae* and one to *Ca. Tenuibacteraceae*. These were differentially associated with different tardigrade species, and fluorescence *in situ* hybridization (FISH) detected bacteria within the ovaries of some tardigrades, suggesting that tardigrade endosymbionts are vertically transmitted (Roberto Guidetti et al. 2020).

A second study surveyed the microbiota of a newly-described tardigrade species, *Paramacrobiotus experimentalis* Kaczmarek, Mioduchowska, Poprawa & Roszkowska, 2020, collected from two samples of moss growing on soil in Madagascar and subsequently raised in laboratory culture for two years before DNA extraction (Kaczmarek et al. 2020). This study again identified differences between the tardigrades’ microbiome and that of their environment and detected evidence of putative endosymbionts of the intracellular groups *Rickettsiales* and *Polynucleobacter*. *Proteobacteria* and *Firmicutes* were the dominant phyla in *Pam. experimentalis*, and 31 operational taxonomic units (OTUs) shared across tardigrade samples were identified as potential core microbiome members for this tardigrade species (Kaczmarek et al. 2020).

A third paper conducted 16S rRNA amplicon sequencing on four tardigrade species: *Hypsibius exemplaris* Gąsiorek, Stec, Morek & Michalczyk, 2018, collected from rotting leaves in a pond in the United Kingdom; *Macrobiotus polypiformis* Roszkowska, Ostrowska, Stec, Janko & Kaczmarek, 2017, collected from moss on a wall in Ecuador; *Paramacrobiotus fairbanksi* Schill, Förster, Dandekar & Wolf, 2010, collected from moss in Antarctica; and *Paramacrobiotus* sp. Guidetti, Schill, Bertolani, Dandekar & Wolf, 2009, collected from moss on a wall, soil, and railroad tracks at two locations in Poland. Of these, all but *Pam. fairbanksi* were subsequently cultured prior to DNA extraction. This study identified *Proteobacteria, Firmicutes*, and *Actinobacteria* as the most abundant phyla in the studied tardigrades, but primarily focused on putative endosymbionts of tardigrades, specifically OTUs assigned to *Rickettsiales* and *Wolbachia*. Members of *Wolbachia* were detected in adult *Pam*. sp. and *Mac. polypiformis*, and *Rickettsiales* were detected in eggs of *Pam. Fairbanksi* as well as adult *Mac. polypiformis* and *Pam*. sp. Neither *Rickettsiales* nor *Wolbachia* were detected in *Hys. exemplaris* or the adult *Pam. fairbanksi* (Mioduchowska et al. 2021).

Most recently, Zawierucha *et al*. sequenced 16S rRNA, ITS1, and 18S rRNA genes to identify bacteria, fungi, and microeukaryotes, respectively, associated with the glacial tardigrade *Cryobiotus klebelsbergi* (Mihelčič, 1959). *C. klebelsbergi* were collected from cryoconite on the surface of Forni Glacier in Italy; DNA was extracted from four samples immediately and from another three after starving for three weeks. The authors found that relative richness of bacteria, fungi, and microeukaryotes was highest in cryoconite, followed by fed tardigrades and finally starved tardigrades. *Polaromonas* sp. was the most abundant bacterium in both fed and starved *C. klebelsbergi*, while *Pseudomonas* sp. and *Ferruginibacter* sp. were the second most abundant bacteria in fed and starved tardigrades, respectively.

16S rRNA gene amplicon sequencing has allowed major advances in understanding of the tardigrade microbiota. However, contamination is an ongoing issue in microbiome studies, especially in low microbial biomass samples such as tardigrades where contaminants can make up a relatively large proportion of all sequence reads and therefore have a disproportionately large impact on results. A minimum standard developed for such studies is the RIDE checklist, which advises researchers to report the methodology used to reduce and assess contamination, to include three types of negative controls (sampling blank, DNA extraction blank, and no-template amplification controls), to determine contamination level by comparing these negative controls to the samples, and to explore contaminants’ impact on results (Eisenhofer et al. 2019). However, while recommended laboratory practices can reduce contamination, they cannot eliminate it. Therefore, *in silico* approaches have been developed to better accomplish the last two steps of the RIDE checklist. For example, the program decontam identifies contaminants based on presence in negative controls and higher frequencies in low-concentration samples. It then removes them from further analysis, dramatically improving the accuracy of results (Davis et al. 2018; Karstens et al. 2019). In this work, we followed the RIDE checklist and utilized decontam for *in silico* contaminant removal.

This study represents the first survey of tardigrade microbiota in North America, as well as the first such survey in an agricultural setting (apple orchards). Rather than focusing on the microbiome of individual tardigrade species, this work is the first to study the microbiome of a full community of tardigrades, hereafter referred to as the tardigrade community microbiome. It is also only the fifth survey of the tardigrade microbiome ever conducted and leverages contamination mitigation methods not used in the previous studies. In addition to identifying putative plant pathogens and endosymbionts associated with tardigrade communities in apple orchards, this study examines whether the tardigrade microbiome differs in four contrasts: (1) across locations, (2) between substrates (moss vs. lichen), (3) between tardigrades and their substrates, and (4) across years.

## 2 Materials and Methods

### 2.1 Moss and Lichen Sample Collection

In summer 2019, lichen samples were collected from apple trees growing in six orchards (Locations 1-6) in Hardin and Franklin counties in north-central Iowa, USA (Fig. S1). One of these (Location 1) had previously been surveyed for tardigrades (Tibbs-Cortes, Tibbs-Cortes, and Miller 2020). One sample of lichen was collected from each tree, and three to five trees were sampled at each location. Moss was also present on the sampled apple trees at Location 2, so a moss sample was collected from three of these trees. In June 2020, additional lichen samples were taken from the trees at Location 1 to enable comparison across years. All moss and lichen samples were placed in individual brown paper bags, which were stored in a cool, dry room to allow the samples to dehydrate. From each of the 2020 lichen samples, five subsamples of 0.25 g were placed in sterile 1.5 mL tubes and frozen at −20 °C for substrate DNA extraction. Table 1 shows a summary of collected samples.

**Table 1.**
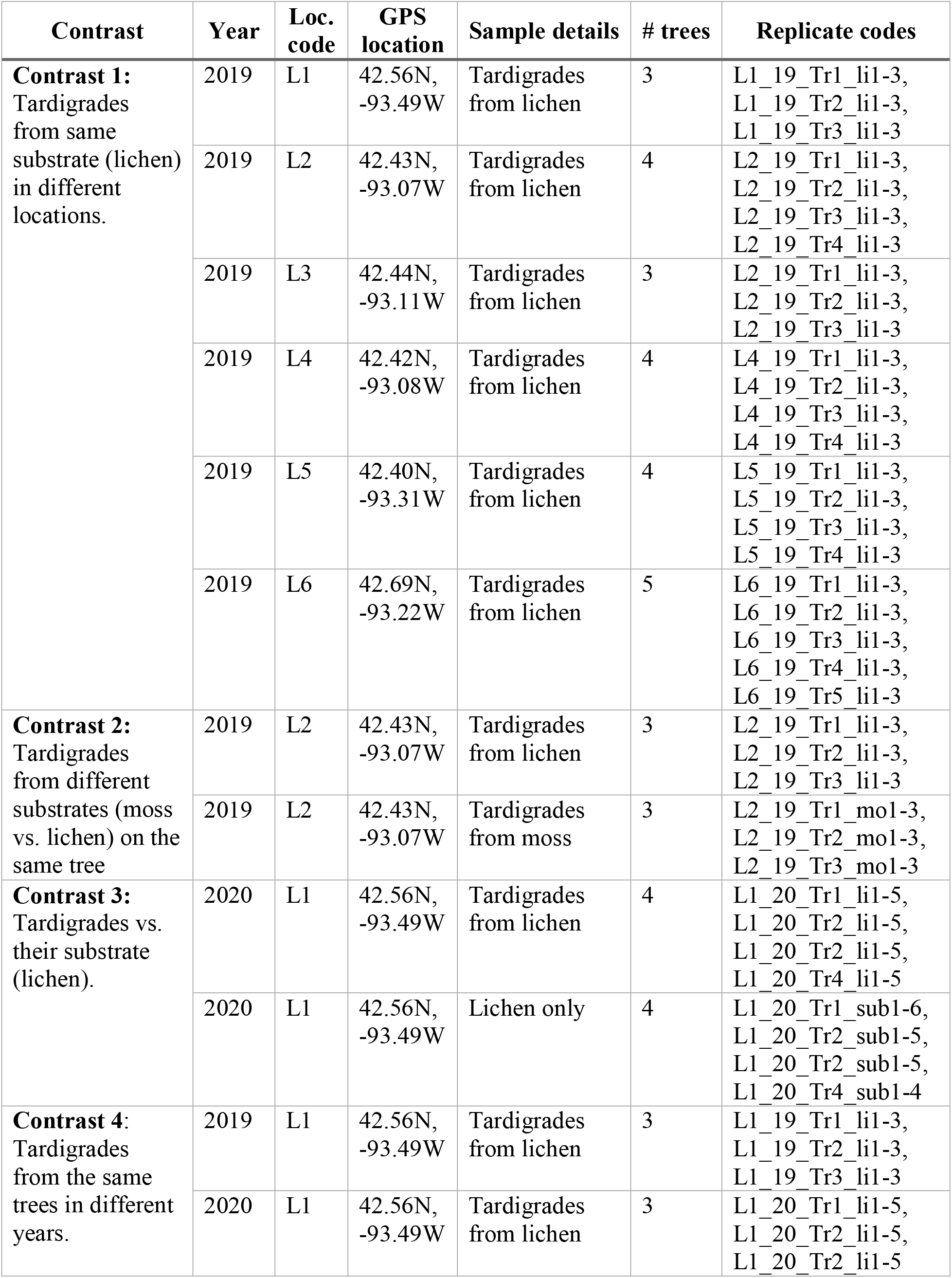
Details of samples used in the experiment. Samples are arranged by contrast; when samples were included in multiple contrasts, these samples appear more than once in the table. “Loc. code” is the location code for a given orchard (e.g., L1 is Location 1) and “# trees” indicates the number of trees sampled at that location for a given contrast. From each sample of moss or lichen, three to six replicates were extracted, identified in the “Replicate codes” column.

### 2.2 Aseptic Technique

16S rRNA gene amplicon sequencing studies focusing on low-biomass samples are prone to biases from external contamination during sample processing, DNA extraction, library preparation, and sequencing. Therefore, all subsequent tardigrade isolation and DNA extraction steps were carried out using barrier pipette tips (Axygen) and in a sterile work area dedicated to the project. A Bunsen burner was used to create a sterile field for all tardigrade isolations and DNA extractions.

### 2.3 Tardigrade Extraction

To extract tardigrades, each moss or lichen sample was soaked in glass-distilled water for a minimum of four hours. Subsamples of this water were then examined under a dissecting microscope, and tardigrades were extracted with Irwin loops (Schram and Davison 2012; Miller 1997). The Irwin loop was disinfected by a flame between each collected tardigrade.

Next, isolated tardigrades were washed by immersion in droplets of PCR-grade water treated with diethyl pyrocarbonate (DEPC). Tardigrades were then transferred to a fresh drop of DEPC-treated water. This washing process was repeated for a total of three washes, and Irwin loops were sterilized between each wash. Three to six replicates of 30 tardigrades each were collected from each substrate sample (identified by replicate codes shown in Table 1) and were then stored in DEPC-treated water at −20 °C.

### 2.4 DNA extraction and sequencing

The DNeasy PowerLyzer PowerSoil Kit (Qiagen) was utilized for DNA extraction. Substrate samples were first ground with a sterilized pestle before being transferred to the bead tubes, while tardigrades were directly transferred to bead tubes. Bead tubes were then transferred to a Bead Mill 24 homogenizer (Fisher Scientific). Tardigrades were homogenized using a single 30 second cycle at 5.00 speed, and substrate samples were homogenized using three 30 second cycles with 10 seconds between cycles at 5.50 speed. Homogenized bead tubes were then centrifuged at 10,000 x *g* (30 seconds for tardigrades and 3 minutes for substrate) before proceeding according to the manufacturer’s instructions; the optional 5 minute incubation at 2 - 8°C was performed during steps 7 and 9. Following elution of DNA with 90 μL of elution buffer, DNA quality and concentration of a 1 μL sample was measured using a NanoDrop spectrometer. Extracted DNA was stored at −20°C.

DNA was loaded onto sterile 96 well plates for library preparation and sequencing. In addition to the tardigrade and substrate samples, three types of controls were included to account for contamination. Six tardigrade processing controls (TPC, equivalent to RIDE sampling blank controls) were created by applying the tardigrade extraction and subsequent DNA extraction protocols to blank samples. Ten DNA processing controls (DPC, equivalent to RIDE DNA extraction blank controls) were created by conducting DNA extraction on 100 μL of DEPC-treated water. Finally, ten wells were loaded with DEPC-treated water to form the library processing controls (LPC, equivalent to RIDE no-template amplification controls). Controls and samples were then submitted for library prep and 16S rRNA gene amplicon sequencing targeting the V4 region at the Iowa State University DNA facility. Library preparation was conducted following the Earth Microbiome Project 16S Illumina amplicon protocol (https://earthmicrobiome.org/protocols-and-standards/16s/) with the following modifications: (1) a single amplification was conducted for each sample rather than in triplicate, (2) PCR purification was conducted using the QIAquick PCR Purification Kit (Qiagen), and (3) all reactions and purification steps were conducted at half volume using a Mantis liquid handler (Formulamatrix) which was cleaned with isopropanol prior to library preparation. Libraries were loaded onto the MiSeq platform at a concentration of ~ 4pM, and paired-end sequencing was conducted at 500 cycles.

### 2.5 Data Analysis

Following sequencing, three paired end samples representing replicates L6_19_Tr2_li2, L6_19_Tr4_li1, and L5_19_Tr3_li3 (Table 1) were removed from the dataset due to poor quality. Raw reads were processed with mothur version 1.43.0. Sequences were screened to remove reads that contained any ambiguities, were shorter than 252 bases, and had homopolymeric sequences greater than eight bases. In total, 1,157,089 reads were removed from the raw dataset of 7,805,248 reads. Screened reads were then aligned against the SILVA alignment version 138, and reads which aligned outside the region covered by 95% of the alignment were removed. The SILVA database was also used to remove 145,663 chimeric sequences and to classify remaining sequences. *De novo* OTU clustering was then conducted at a 99% similarity threshold.

R version 4.0.3 running packages decontam (Davis et al. 2018), phyloseq (McMurdie and Holmes 2013), and corncob (Martin, Witten, and Willis 2020), as well as a more efficient implementation of DivNet known as divnet-rs (https://github.com/mooreryan/divnet-rs) running in Rust, were used for subsequent analyses. Using decontam, contaminant OTUs were identified and removed based on their relative prevalence in control vs. true samples (prevalence method, threshold 0.25) (Davis et al. 2018). Next, OTUs with fewer than 10 reads in experimental samples were removed. From these data, alpha diversity parameters (Shannon and Simpson) were calculated using DivNet and divnet-rs (Willis and Martin 2020). Relative abundance, differential abundance, and differential variability of taxa were calculated in corncob using a beta-binomial model (Martin, Witten, and Willis 2020). Differences were declared significant when False Discovery Rate (FDR)-corrected *P* values (Benjamini and Hochberg 1995) were less than 0.05. Principal Coordinates Analysis (PCoA) was conducted in phyloseq using the default Bray-Curtis distance.

### 2.6 Identification of unclassified putative plant pathogens and endosymbionts

In the cases where OTUs of interest were not classified by mothur to the genus level, BLAST and RDP Classifier (Camacho et al. 2009; Wang et al. 2007) results were used to provide additional information about taxonomic classification. First, the mothur command “get.oturep” was used to generate a FASTA file containing the representative sequence for each OTU; OTUs with fewer than 10 reads in experimental samples were removed. BLAST analysis was performed using BLAST+ v2.11.0. The NCBI 16S RefSeq collection (representing 22,061 taxa) (O’Leary et al. 2016) was downloaded and converted into a BLAST database using the “makeblastdb” command. The “blastn” command was then run against this database using the representative sequence FASTA as the query. Results with the 15 lowest E-values were kept for each OTU. The representative sequence FASTA was also entered into the RDP Classifier web tool version 2.11 using 16S rRNA training set 18 (https://rdp.cme.msu.edu/classifier/classifier.jsp), and the assignment detail for all OTUs was downloaded.

### 2.7 Data and Code Availability

Raw sequencing files are deposited at the Sequence Read Archive. Mothur output and code used for analysis is available at https://github.com/LTibbs/tardigrade_microbiome.

## 3 Results

In total, 118 DNA samples and 26 controls were sequenced. The DNA samples consisted of 20 from lichen as well as 89 and 9 from tardigrades extracted from lichen and moss, respectively, collected from a total of 23 different apple trees in six Iowa orchards. The controls consisted of 6 TPCs, 10 DPCs, and 10 LPCs. From these sequences, 248,493 OTUs were identified by mothur. The decontam package identified and removed 986 OTUs as contaminants. Of the remaining OTUs, 235,652 were removed because they were represented by fewer than ten reads in the experimental samples, leaving 11,855 OTUs for further analysis.

Mothur classification and decontam scores for all OTUs with more than 10 reads in experimental samples are shown in Table S1. BLAST results and RDP Classifier results for these OTUs can be found in Tables S2 and S3, respectively. Relative abundance of OTUs by sample and by contrast are provided in Table S4 and Tables S5-S8, respectively; significantly differentially abundant and variable phyla, genera, and OTUs across contrast levels are presented in Table S9. Overall, the five most abundant phyla were *Proteobacteria, Bacteroidota, Actinobacteriota, Firmicutes*, and *Acidobacteriota* (Fig. S2), while the three most abundant genera were *Pseudomonas, Bradyrhizobium*, and an unclassified *Enterobacteriaceae* (Fig. S3). From the PCoA of all samples, the first principal coordinate clearly separates substrate samples from tardigrade samples, while the second coordinate tends to separate the 2020 from the 2019 samples. Samples from different locations and from moss and lichen are not clearly separated by the first two coordinates (Fig. S4).

### 3.1 Contrast 1: Location

The tardigrade community microbiome differed significantly across locations, as shown by the Simpson and Shannon indices, which differed significantly in most pairwise comparisons of locations (Table 2). Across locations, 13 phyla and 44 genera were both significantly differentially abundant and significantly differentially variable. Sixteen OTUs were significantly differentially abundant only, four OTUs were significantly differentially variable only, and three OTUs were both significantly differentially abundant and variable (Table S9). These identified differential taxa included the aforementioned top five phyla (*Proteobacteria, Bacteroidota, Actinobacteriota, Firmicutes*, and *Acidobacteriota*) and top three genera (*Pseudomonas, Bradyrhizobium*, and unclassified *Enterobacteriaceae*) from the experiment as a whole. Despite these differences, the locations clustered together in the PCoA (Fig. 1).

**Table 2.** Alpha diversity measures Shannon and Simpson estimated for each level of each contrast of interest. Within each contrast and diversity measure, estimates that are significantly different from one another (Benjamini-Hochberg corrected *P* value <0.05) share no letters.

| Contrast | Level of Contrast | Shannon | Simpson |
| --- | --- | --- | --- |
| 1: Location | Location 1 | 1.570<sup>a</sup> | 0.437<sup>a</sup> |
|  | Location 2 | 3.545<sup>b</sup> | 0.244<sup>b</sup> |
|  | Location 3 | 0.970<sup>c</sup> | 0.779<sup>c</sup> |
|  | Location 4 | 2.093<sup>d</sup> | 0.466<sup>a</sup> |
|  | Location 5 | 1.605<sup>a</sup> | 0.683<sup>d</sup> |
|  | Location 6 | 3.830<sup>b</sup> | 0.255<sup>b</sup> |
| 2: Moss vs. Lichen | Moss | 2.929<sup>a</sup> | 0.175<sup>a</sup> |
|  | Lichen | 2.926<sup>a</sup> | 0.205<sup>a</sup> |
| 3: Tardigrades vs. Substrate | Tardigrade | 2.895<sup>a</sup> | 0.279<sup>a</sup> |
|  | Substrate | 6.673<sup>b</sup> | 0.004<sup>b</sup> |
| 4: Year | 2019 | 1.448<sup>a</sup> | 0.447<sup>a</sup> |
|  | 2020 | 1.566<sup>a</sup> | 0.365<sup>b</sup> |

**Figure 1.**
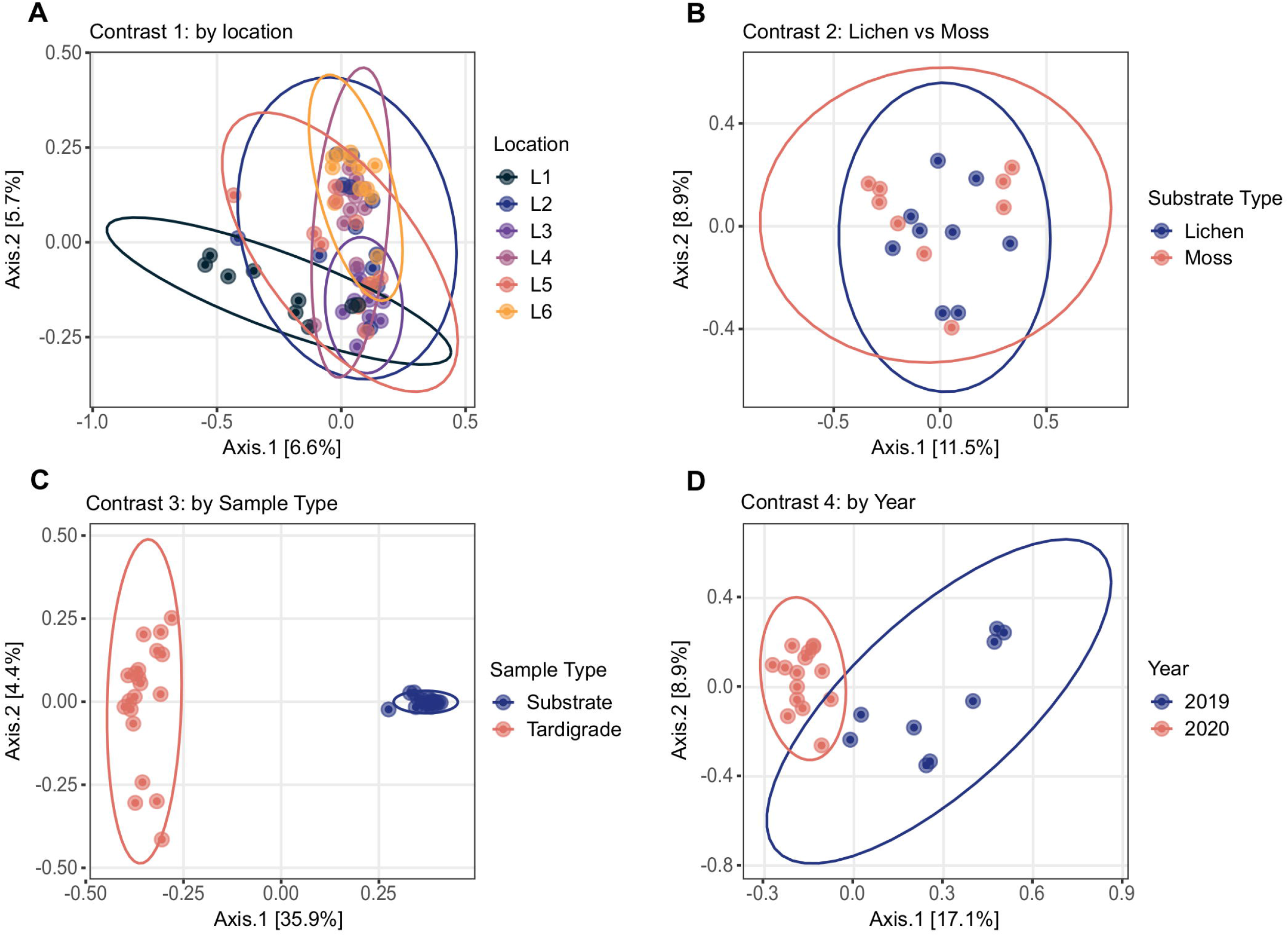
Principal Coordinates Analyses conducted by contrast based on Bray-Curtis distance. Location names (Location 1 through Location 6) are abbreviated as L1 through L6. **(A)** Tardigrade samples from different locations overlap one another, as do **(B)** tardigrades isolated from lichen and moss. However, **(C)** tardigrade and substrate samples are clearly separated, and **(D)** 2019 and 2020 tardigrade samples are mostly separated.

### 3.2 Contrast 2: Moss vs. Lichen

The community microbiome of tardigrades extracted from moss did not differ significantly in alpha diversity from that of tardigrades extracted from lichen as measured by the Shannon and Simpson indices (Table 2). PCoA further demonstrates that the overall microbial community did not differ by substrate type (Fig. 1). However, between tardigrades collected from moss and those from lichen, five phyla and 11 genera were both significantly differentially abundant and significantly differentially variable, while three OTUs were significantly differentially abundant only (Table S9). These included the common phyla *Firmicutes* and *Bacteroidota*; of these, *Firmicutes* were more abundant in moss-associated and *Bacteroidota* in lichen-associated tardigrades (Table S10, Fig. 2).

**Figure 2.**
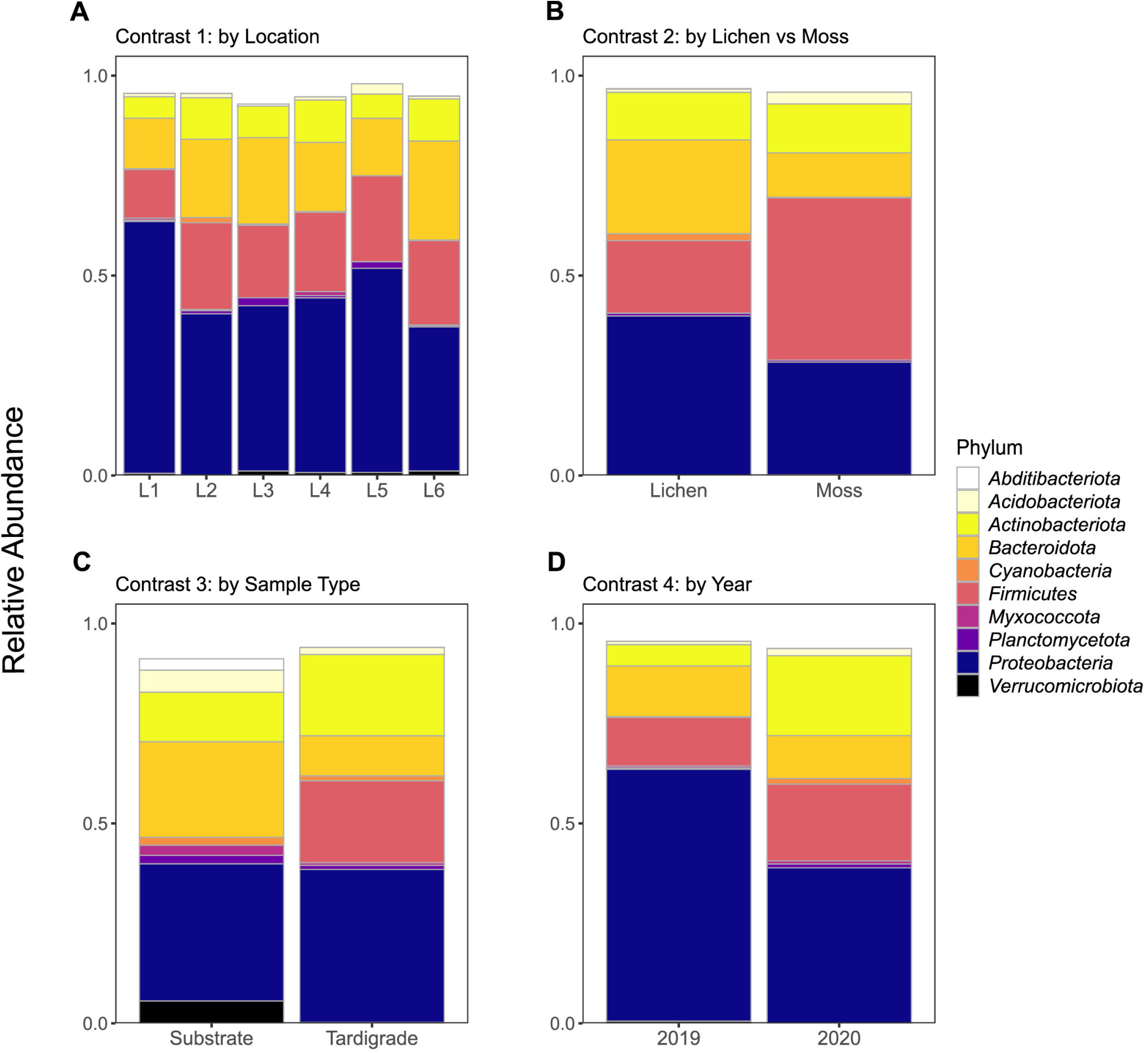
Relative abundance of top 10 identifiable phyla shown across all contrasts. Location names (Location 1 through Location 6) are abbreviated as L1 through L6.

### 3.3 Contrast 3: Tardigrades vs. Substrate

The microbiota of tardigrades was significantly less diverse than that of their lichen substrate, as measured by both Shannon and Simpson indices (Table 2); the tardigrade and substrate samples also formed distinct clusters as shown by PCoA (Fig. 1). Between tardigrades and their substrate, 17 phyla, 181 genera, and 101 OTUs were significantly differentially abundant and variable, while 308 OTUs were significantly differentially abundant only and 124 OTUs were significantly differentially variable only (Table S9). These differential taxa included four of the top five phyla (all except *Proteobacteria*) and all three of the three most abundant genera from the experiment as a whole. Remarkably, the relative abundance of *Firmicutes* was nearly a thousand times higher in the tardigrades (20.5%) than in their substrate (0.021%) (Table S10, Fig. 2).

### 3.4 Contrast 4: Year

From 2019 to 2020, the tardigrade community microbiome increased in diversity as measured by the Simpson index, though no significant difference was found between the Shannon indices (Table 2). The two years also formed mostly distinct clusters in the PCoA (Fig. 1). Between the two years, 44 genera and one OTU were significantly differentially variable and abundant, while two phyla and 26 OTUs were significantly differentially abundant only (Table S9). These differential taxa included two of the five most common phyla (*Proteobacteria* and *Actinobacteriota*) and two of the three most common genera (*Pseudomonas* and unclassified *Enterobacteriaceae*). The unclassified *Enterobacteriaceae* had a particularly large change in relative abundance, decreasing more than 160-fold from 10.1% in 2019 to 0.062% in 2020 (Table S11, Fig. 3).

**Figure 3.**
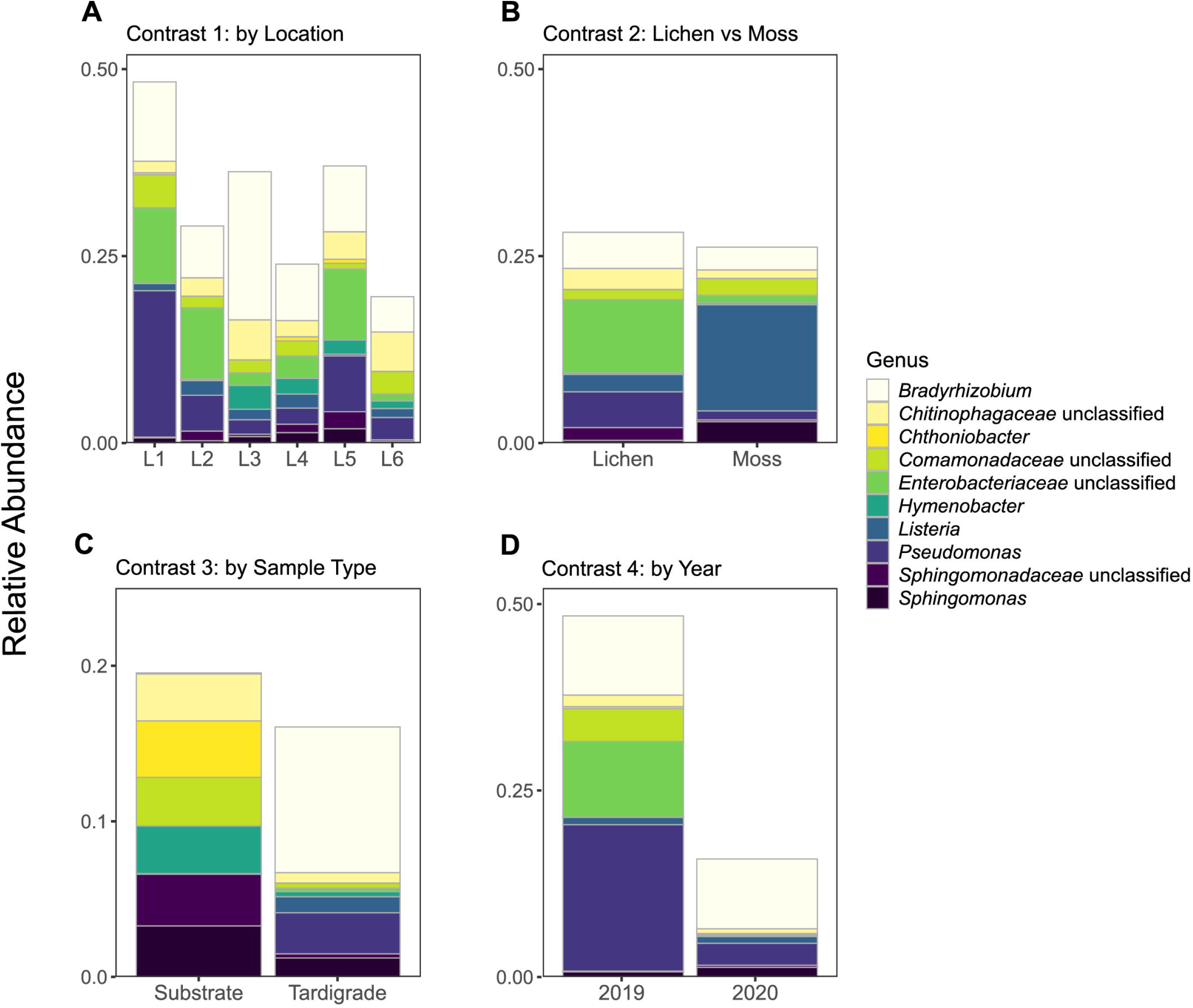
Relative abundance of top 10 genera (identifiable at least to family level) shown across all contrasts. Location names (Location 1 through Location 6) are abbreviated as L1 through L6.

## 4 Discussion

### 4.1 Tardigrade Community

This study examined the microbiota of the full tardigrade community from a particular substrate sample, in contrast to previous surveys that studied isolated species, often from laboratory cultures (Vecchi et al. 2018; Kaczmarek et al. 2020; Mioduchowska et al. 2021; Zawierucha et al. 2022). It is of course desirable to identify species-specific microbiota, as Vecchi et al. (2018) found that tardigrade-associated bacteria varied among tardigrade species. However, while it is occasionally possible in samples containing only a few tardigrade species to extract members of each for study (Vecchi et al. 2018), many environmental samples contain numerous species, including cryptic species that may be difficult or impossible to distinguish without molecular, life cycle, or other data (Roberto Guidetti et al. 2016; Cesari et al. 2013). For example, in the current survey, at least three tardigrade genera (*Milnesium, Ramazzottius*, and *Paramacrobiotus*) were observed, but all would require additional morphometric or egg observations to identify to species level (Michalczyk et al. 2012; R. Guidetti et al. 2009; Kinchin 1996; Binda 1987). Consequently, laboratory culturing is usually required for identification of tardigrade species, but Vecchi et al. (2018) found that culturing significantly affects the tardigrade microbiome. Therefore, the current study has the unique advantage of better reflecting the tardigrade community microbiome in its natural state compared to studies that focus on cultured tardigrades.

While tardigrade species were not identified in this study, a December 2015 collection effort at Location 1 provides information on tardigrade diversity in the area. From lichen growing on some of the same apple trees used in the current study, the previous study identified *Milnesium* cf. *barbadosense* Meyer and Hinton 2012; *Mil. burgessi* Schlabach, Donaldson, Hobelman, Miller, and Lowman, 2018; *Mil. swansoni* Young, Chappell, Miller, and Lowman, 2016; and *Pam. (A.) tonollii* (Ramazzotti, 1956), as well as members of *Milnesium* Doyère, 1840; *Ramazzottius* Binda and Pilato, 1986; *Paramacrobiotus* Guidetti, Schill, Bertolani, Dandekar and Wolf, 2009; and Macrobiotidae Thulin, 1928 not identifiable to species (Tibbs-Cortes, Tibbs-Cortes, and Miller 2020). While tardigrade communities are dynamic across both time (Schuster and Greven 2007, 2013) and space (Meyer 2008, 2006), the dominant species present in tardigrade communities in a given area can remain remarkably stable across years (Schuster and Greven 2007; Nelson and McGlothlin 1996). This suggests that species information from the 2015 survey may be relevant to the current study.

### 4.2 Reducing Effects of Contamination

All tardigrade microbiome surveys, including the current study, employed laboratory technique to reduce contamination by washing the tardigrades in sterile water before DNA extraction (Vecchi et al. 2018; Mioduchowska et al. 2021; Kaczmarek et al. 2020; Zawierucha et al. 2022). Working in a sterile environment further decreases contamination; therefore, Zawierucha *et al*. extracted tardigrades from substrate in a sterile environment (laminar flow chamber) (2022), and we worked in a sterile field created by a Bunsen burner throughout the experiment. Three previous studies included one type each of negative controls recommended by the RIDE standards for low biomass studies (Eisenhofer et al. 2019) (DNA extraction blank in (Mioduchowska et al. 2021), sampling blank in (Kaczmarek et al. 2020), and no-template amplification control in (Zawierucha et al. 2022)). Kaczmarek *et al*. and Mioduchowska *et al*. did not sequence the negative controls; instead, they performed PCR amplification of these controls and determined that no contamination was present because no bands were visible (Kaczmarek et al. 2020; Mioduchowska et al. 2021). However, samples without visible bands from PCR can generate sequencing reads (Davis et al. 2018) and would not detect contaminants introduced during library preparation or sequencing steps. Zawierucha *et al*. removed all OTUs present in the no-template amplification control from analysis (2022). However, low levels of true sequences, especially from high-abundance OTUs, are often present in negative controls due to cross-contamination of samples; these biologically important OTUs would therefore be removed from the analysis (Davis et al. 2018; Karstens et al. 2019). In our study, we included and sequenced all three RIDE-recommended types of negative controls.

No previous survey of the tardigrade microbiota has employed model-based *in silico* contaminant identification and removal, which we accomplished using the decontam package. Decontam removed 986 OTUs as contaminants, including five that would otherwise have been in the top ten OTUs in the study by read count. Of course, further improvements are always possible. Contaminants are expected to differ in prevalence among negative control types depending on their point of introduction, but current *in silico* contamination removal methods treat all negative controls identically (Davis et al. 2018). Future development of a method that leverages the unique information provided by each type of negative control would therefore be desirable. However, by working under a flame, including and sequencing all recommended types of negative controls, and leveraging *in silico* contaminant identification and removal, we have produced what we expect to be the tardigrade microbiome survey least affected by contamination to date.

### 4.3 Tardigrade Community Microbiome by Contrast

We investigated the tardigrade community microbiome across four contrasts. First, we determined that the tardigrade community microbiome varied significantly in structure across locations (Table 2). Vecchi *et al*. (2018) found that an average of 15.4% of the microbial OTUs in a tardigrade collected from moss or lichen originate from its substrate. Therefore, known impacts of geographical location on microbial communities (Baldrian 2017; Coller et al. 2019) could have resulted in different microbial communities present in each location to inoculate the tardigrades. Additionally, as the tardigrade microbiota is species-specific (Vecchi et al. 2018), the differences in microbial communities observed across locations may reflect spatial variation in tardigrade communities’ species composition (Meyer 2008). Of course, these explanations are not mutually exclusive and could both play a role in shaping distinct tardigrade community microbiomes across locations. Vecchi *et al*. surveyed tardigrades of the same species collected from different locations, but they did not test for differences in the microbiome across locations. However, they did identify an OTU in the genus *Luteolibacter* that was significantly associated with *Ram. oberhaeuseri* collected from a location at 34 meters above sea level but not from another location 797 meters above sea level (Vecchi et al. 2018), suggesting that future work may detect differences in the microbiota of the same tardigrade species across locations.

In contrast two, the community microbiome of tardigrades collected from lichen was compared with that of tardigrades collected from moss on the same trees. While a few taxa were significantly differentially abundant and variable across substrates, the two substrates did not differ in alpha diversity and were not separated by PCoA (Fig. 1). This similarity in the tardigrade community microbiome was initially surprising, as previous literature has demonstrated significant differences between the microbiota of moss and lichen even on the same tree (Aschenbrenner et al. 2017). However, this similarity across substrates could be due to the presence of similar tardigrade species, as previous studies have failed to demonstrate significant differences between tardigrade communities found in moss and lichen (Young and Clifton 2015; Nelson, Bartels, and Fegley 2020). In future surveys, it would be interesting to compare the microbiota of tardigrades from additional substrate types (e.g., soil) and to determine if this similarity in tardigrade microbiome across substrates persists at the species as well as the community level.

Results of contrast three demonstrate that the tardigrade community microbiome is distinct from and significantly less diverse than that of its lichen substrate (Table 2, Fig. 1). This result agrees with previous studies that found relatively higher diversity in substrates than in their resident Ecdysozoans, including wild tardigrades collected from moss and lichen (Vecchi et al. 2018), cultured tardigrades (Kaczmarek et al. 2020), and the nematodes *Meloidogyne hapla* (Adam et al. 2014) and *Caenorhabditis elegans* (Johnke, Dirksen, and Schulenburg 2020). This study is the first to demonstrate this trend at the tardigrade community rather than species level. Vecchi et al. (2018) suggested that the lower microbial diversity in tardigrades with respect to their substrates may be due to the small size of tardigrades limiting the biomass and therefore the diversity of their microbiome (small host hypothesis) and/or to selectiveness of tardigrades inhibiting growth of some bacterial species and promoting growth of others (selective host hypothesis). Supporting the selective host hypothesis, earlier work found that tardigrades could be successfully inoculated with some bacteria (*Xanthomonas*) but not others (*Serratia*) (Krantz, Benoit, and Beasley 1999). It is possible that some bacteria have co-evolved with tardigrades, becoming permanent residents of the gastrointestinal tract or cuticle, a hypothesis that has been suggested for the Ecdysozoan *C. elegans* (F. Zhang et al. 2017). The life cycle of tardigrades poses a unique selective pressure on any permanent residents of the microbiota, as these organisms would also have to survive within the tardigrade during cryptobiosis. This would be especially true for the obligate endosymbiotic taxa *Rickettsiales* and *Polynucleobacter* previously observed in tardigrades (Vecchi et al. 2018; Roberto Guidetti et al. 2020; Kaczmarek et al. 2020; Mioduchowska et al. 2021), as well as for the *Rickettsia* identified in the current study (see below).

Contrast four determined that the tardigrade community microbiome is temporally dynamic, changing significantly on the same trees from 2019 to 2020 (Table 2, Fig. 1). Again, this may be due to changes in habitat microbiome, as microbiota of other substrates (e.g., soil and litter) are known to vary across years due to changing environmental factors such as nutrient availability (Martinović et al. 2021). This variation may also be due to temporal changes in the tardigrade community composition; although tardigrade species present may remain consistent in a location over years, their relative abundances shift in part due to changes in rainfall, humidity, and temperature (Schuster and Greven 2007). This temporal variability raises important implications for future studies of the tardigrade community microbiome. For example, the relative abundance of putative phytopathogens differed significantly across years (Table S9). Future work could identify temporal variables affecting the ability of tardigrades to act as potential reservoirs of phytopathogens and other bacteria. We also encourage further studies of the tardigrade microbiome to account for temporal changes and to investigate this variation with additional time points to increase resolution.

### 4.4 Tardigrade-Associated Taxa

In this study, the five most abundant phyla were *Proteobacteria, Firmicutes, Bacteroidota, Actinobacteria*, and *Acidobacteria* (Table S10). All of these except *Acidobacteria* were previously reported as highly abundant in at least two of the three previous tardigrade microbiome surveys that presented results at a phylum level, with *Proteobacteria* identified as the most abundant phylum in all cases (Vecchi et al. 2018; Kaczmarek et al. 2020; Mioduchowska et al. 2021). Combined, the tardigrades in these studies represent a diverse set of species, including wild and laboratory-reared specimens isolated from multiple continents, suggesting that the predominance of these phyla is broadly characteristic of the microbiome of Tardigrada, regardless of species or location. These phyla, especially *Proteobacteria*, are also dominant in the microbiomes of other Ecdysozoans, including soil nematodes (Dirksen et al. 2016; Elhady et al. 2017; Adam et al. 2014), marine nematodes (Arcos et al. 2021), and insects (Colman, Toolson, and Takacs-Vesbach 2012; Engel and Moran 2013). The tardigrade microbiota therefore appears similar to that of other Ecdysozoans at the phylum level.

A number of OTUs significantly more abundant in tardigrades than in their substrate in this study belong to taxa previously identified in the tardigrade microbiome. These include members of *Enhydrobacter* (Vecchi et al. 2018), *Enterobacteriaceae* (Mioduchowska et al. 2021; Kaczmarek et al. 2020), and *Acinetobacter* (Vecchi et al. 2018; Mioduchowska et al. 2021) (Table S9). In this study, OTU 22 was classified as *Enhydrobacter*, and its abundance in the tardigrade population varied over time, increasing significantly from 2019 (0.21%) to 2020 (2.1%) (Table S8, Table S9). It is possible that *Enhydrobacter* is common to Ecdysozoan microbiomes, as it is also an abundant taxon in the gut contents of larval wood wasps (J. Li et al. 2021) and nematodes (Adam et al. 2014). Members of *Enterobacteriaceae* included OTUs 2 and 20. OTU 20 was further identified as a member of the *Escherichia/Shigella* complex, but OTU 2 could not be classified to the genus level (Table S2, Table S3). OTU 2 showed significant temporal variation, decreasing in relative abundance from 10.0% to 0.062% from 2019 to 2020 (Table S8, Table S9). *Enterobacteriaceae* is also highly represented in the gut microbiota of insects (Moro et al. 2021; Hernández-García et al. 2017) and nematodes (Zhou et al. 2022; Zimmermann et al. 2020). This suggests that *Enterobacteriaceae* may be residents of the tardigrade digestive tract. Finally, OTU 16 was a member of *Acinetobacter* that increased significantly in abundance from 2019 (0.000052%) to 2020 (2.3%) and was one of the three OTUs significantly differentially abundant across substrate type (Table S8, Table S9). *Acinetobacter* is associated with the cuticle of the nematodes *M. hapla, M. incognita*, and *Pratylenchus penetrans* (Adam et al. 2014; Elhady et al. 2017), suggesting that it may also be associated with the cuticle of tardigrades.

However, many of the tardigrade-associated taxa observed in this study have not been previously reported in the tardigrade microbiome. In fact, the most abundant OTU across all samples in this study (OTU 1) was a member of the genus *Bradyrhizobium*, which was not previously reported from the tardigrade microbiome. This OTU was also spatially dynamic, differing significantly in abundance across locations (Table S9). *Bradyrhizobium* has been previously observed in the microbiota of plant pathogenic nematodes (Eberlein et al. 2016; Adam et al. 2014) and leaf hoppers (Horgan et al. 2019). This genus has also been found in the lichen microbiome (Bates et al. 2011; Erlacher et al. 2015; Graham et al. 2018), perhaps indicating that tardigrades acquire this bacterium from their habitat. Another tardigrade-associated genus, *Micrococcus*, was differentially abundant across both locations and years (Table S9). This genus has been reported from the cuticles of soil nematodes (Adam et al. 2014) as well as fish parasitic nematodes (Arcos et al. 2021), suggesting that *Micrococcus* may be associated with the tardigrade cuticle. Another notable tardigrade-associated genus in this study was *Nakamurella*, represented primarily by OTU 33, which showed differential abundance across locations (Table S9). *Nakamurella intestinalis* has been isolated from the feces of another Ecdysozoan, the katydid *Pseudorynchus japonicus* (Kim et al. 2017). *N. endophytica* and *N. flava* were identified as endophytes of mangroves and mint, respectively (Yan et al. 2020; Tuo et al. 2016), and *N. albus* and *N. leprariae* were originally discovered in lichens (Jiang et al. 2020; An et al. 2021). This suggests that tardigrades could obtain endophytic or lichen-dwelling *Nakamurella* from their habitat.

### 4.5 Putative Endosymbionts

Our survey corroborates previous observations of putative endosymbionts of the obligate intracellular order *Rickettsiales* associated with tardigrades. Three of the four previous surveys of the tardigrade microbiome have detected OTUs of this order (Vecchi et al. 2018; Roberto Guidetti et al. 2020; Kaczmarek et al. 2020; Mioduchowska et al. 2021); in addition to unclassified *Rickettsiales*, these OTUs included members of *Wolbachia* (Mioduchowska et al. 2021), *Anaplasmataceae*, and *Ca. Tenuibacteraceae* (Roberto Guidetti et al. 2020). Kaczmarek *et al*. also detected the obligate intracellular genus *Polynucleobacter* (2020). In the current survey, we identified two *Rickettsiales* OTUs. Of these, one was classified by mothur as *Wolbachia* (OTU 3606), and the other was further classified as *Rickettsia* (OTU 180) by BLAST and RDP analysis (Table S2, Table S3). The relative abundance of OTU 180 was significantly higher in tardigrades (0.88%) than in their lichen substrate (0.0012%) (Table S7) as well as significantly higher in 2020 (1.0%) than in 2019 (0.00026%) (Table S8, Table S9). OTU 3606 was numerically more abundant in tardigrades (0.030%) than substrate (0.0000000000051%), though this difference was not statistically significant (Table S7). Taken together, the intracellular nature of *Rickettsiales* and the higher abundance in tardigrades suggests that OTUs 180 and 3606 are endosymbionts of tardigrades.

The presence of endosymbionts may have implications for tardigrade reproduction and evolution, as members of *Rickettsia* and *Wolbachia* are known to manipulate host reproduction in other Ecdysozoans. *Wolbachia* is well-known for causing parthenogenesis in nematodes and arthropods, as well as feminization of males, cytoplasmic incompatibility, and male-killing (Werren, Baldo, and Clark 2008; Kraaijeveld et al. 2011; Correa and Ballard 2016; Kajtoch and Kotásková 2018). Similarly, *Rickettsia* can induce parthenogenesis (Hagimori et al. 2006; Giorgini et al. 2010) and male-killing (Lawson et al. 2001) in arthropods. Parthenogenesis is common in tardigrades (Bertolani 2001; Guil et al. 2022). Further investigation is necessary to determine if this is due to reproductive manipulators such as *Rickettsia* and *Wolbachia*. Future analysis could follow the example of Guidetti *et al*. by incorporating FISH to confirm the presence of these and other endosymbionts within tardigrade tissues (2020).

### 4.6 Putative Phytopathogens

Our analysis also aimed to determine whether wild tardigrades living in apple orchards harbor phytopathogenic bacteria, and in fact, the second most abundant genus overall found in this survey was *Pseudomonas*, which contains more than twenty known plant pathogens (Höfte and De Vos 2006). *Pseudomonas* was significantly associated with tardigrades (relative abundance in tardigrades and substrate of 2.7% and 0.051%, respectively) (Table S11) and was spatially and temporally dynamic in the tardigrade community microbiome, as relative abundance of *Pseudomonas* decreased significantly from 2019 (19.6%) to 2020 (3.0%) and differed significantly across locations (Table S9, Table S11). *Pseudomonas* was also detected in all four of the previous surveys of the tardigrade microbiome (Vecchi et al. 2018; Mioduchowska et al. 2021; Kaczmarek et al. 2020; Zawierucha et al. 2022), and Vecchi *et al*. identified it as part of the core tardigrade microbiome (2018). *Pseudomonas* is also present in the microbiota of soil nematodes (Adam et al. 2014; Dirksen et al. 2016; Zimmermann et al. 2020) and insects (Hernández-García et al. 2017; Horgan et al. 2019; Xue et al. 2021). Notably, other Ecdysozoans (insects) act as vectors of *P. syringae* (Stavrinides, McCloskey, and Ochman 2009; Donati et al. 2017), which is one of the most agriculturally damaging *Pseudomonas* species (Höfte and De Vos 2006; Xin, Kvitko, and He 2018). However, *Pseudomonas* is very diverse, containing many non-pathogenic species (Silby et al. 2011; Passera et al. 2019). In fact, some *Pseudomonas* isolates from wild *C. elegans* confer resistance to fungal pathogens in their hosts (Dirksen et al. 2016), raising the possibility that *Pseudomonas* could be similarly beneficial to tardigrades.

Two additional putative phytopathogens were significantly more abundant in tardigrades than their substrate. The first, OTU 261, was identified by mothur as a member of *Ralstonia*, a genus containing the phytopathogenic *R. solanacearum* complex. In addition to being found at significantly higher abundance in tardigrades (0.018%) than their substrate (.00017%) (Table S7), OTU 261 was temporally dynamic, decreasing significantly from 2019 (0.16%) to 2020 (0.00068%) (Table S8, Table S9). *Ralstonia* has been previously observed in the tardigrade *Pam. fairbanksi* (Mioduchowska et al. 2021) and in nematodes (Elhady et al. 2017; Eberlein et al. 2016). The *R. solanacearum* complex causes major yield losses in food crops including tomatoes, bananas, and potatoes (Yuliar, Nion, and Toyota 2015; Paudel et al. 2020). Two notable members of this complex are spread by insect vectors; the cercopoids *Hindola fulva* and *H. strata* act as vectors of *R. syzygii*, while the Blood Disease Bacterium is spread nonspecifically by pollinators (Eden-Green et al. 1992; Remenant et al. 2011).

The second, OTU 208, was classified by BLAST and RDP analysis to the *Erwinia/Pantoea* cluster (Table S2 Table S3), which includes a number of economically important phytopathogens (Kido et al. 2008; Y. Zhang and Qiu 2015; Dutkiewicz et al. 2016; Shapiro et al. 2016). *E. amylovora* is of particular note as it causes fire blight in apple trees (Aćimović et al. 2015). This OTU had a significantly higher relative abundance of 0.046% in tardigrades compared to 0.0044% in their substrate (Table S7, Table S9). While neither *Erwinia* nor *Pantoea* have previously been identified in tardigrades, *Erwinia* has been found in arthropods (Xue et al. 2021) and nematodes (Eberlein et al. 2016). Additionally, multiple phytopathogens in *Pantoea* and *Erwinia* are transmitted by insect vectors (Dutkiewicz et al. 2016; Ordax et al. 2015; Sasu et al. 2010; Basset et al. 2000; Walterson and Stavrinides 2015). However, it is also possible that OTU 208 represents a symbiont in tardigrades, as *Erwinia* also includes the olive fly obligate gut symbiont *Candidatus Erwinia dacicola* (Blow et al. 2020).

Additional putative plant pathogens were observed at lower abundances in the tardigrade community microbiome and were not significantly more abundant in tardigrades than their substrate. These include another *Ralstonia* (OTU 1556) and OTU 1620, which was classified as *Pectobacterium* by BLAST and RDP (Table S2, Table S3). Members of *Pectobacterium* cause soft rot diseases in economically important plants, and some strains are capable of infecting multiple plant species (Ma et al. 2007; X. Li et al. 2020). Additionally, prompted by previous observation of the tardigrade *Mac. hufelandi* acting as a vector of the plant pathogen *Xanthomonas campestris* (Benoit et al. 2000), we searched the tardigrade community microbiome for members of *Xanthomonas*. OTUs 10,409 and 12,281 were classified as *Xanthomonas* (OTU 10,409 by BLAST and RDP analysis), but were both at extremely low abundance (Table S4).

In summary, we observed the presence of multiple putative plant pathogens in the community microbiome of tardigrades isolated from apple orchards. Tardigrades could act as vectors or reservoirs of these putative pathogens, a possibility raised by the previous observation of *Mac. hufelandi* as a vector of *X. campestris* (Benoit et al. 2000). However, a major limitation of this study is the use of only 16S rRNA amplicon sequencing. Because multiple marker genes are required to distinguish among species within *Pseudomonas, Ralstonia, Erwinia*, and *Pantoea* (Y. Zhang and Qiu 2015; Paudel et al. 2020; Palmer et al. 2017; Gomila et al. 2015; Saati-Santamaría et al. 2021), we were unable to identify OTUs in our study to species level. Therefore, we are unable to determine whether the identified OTUs in plant pathogenic genera are themselves phytopathogens. We encourage future analyses of tardigrade-associated bacteria in these groups through techniques such as metagenome sequencing and multilocus sequence typing to clarify this point.

## 5 Conclusion

This study is the first microbiome analysis of wild tardigrade populations in an agricultural setting and is also the first microbiome study assessing North American tardigrades. Our methods reduced the effects of contamination compared to other tardigrade microbiome studies by including aseptic technique, all three recommended control types, and *in silico* contaminant removal. We found that the tardigrade community microbiome is distinct from the substrate microbiota and varies across location and time. In addition to identifying putative endosymbionts, we also observed multiple tardigrade-associated taxa that may represent phytopathogens. The results of this study both increase our knowledge of the tardigrade microbiome and prompt new avenues of research.

## 6 Author Contributions

LTC: Conceptualization, Software, Formal Analysis, Investigation, Data Curation, Writing – Original Draft and Review & Editing, Visualization. BTC: Conceptualization, Investigation, Data Curation, Writing – Original Draft and Review & Editing, Visualization. SSE: Methodology, Resources, Writing - Review & Editing

## 7 Acknowledgements

We wish to thank the owners of the studied apple orchards, including Dennise Smith, Roland Newby, Koenigs’ Acres Farm, and others who wish to remain anonymous. We also thank Lucas Koester and Chiron Anderson for providing example phyloseq code and tutorials as well as useful input on the project. We thank the Iowa State University (ISU) DNA Facility for their input on sequencing, the ISU Department of Ecology, Evolution, and Organismal Biology for use of microscopes, the Iowa Geological Survey for shapefiles used to produce the collection map, and the ISU Plant Sciences Institute for funding. Laura and Bienvenido Tibbs-Cortes are supported by the National Science Foundation Graduate Research Fellowship Program (NSF GRFP Grant No. 1744592).

## Figure Legends

**Figure S1.** Collection locations map. The map of Iowa, USA shows the sampled counties outlined in red. The inset shows collection sites within Hardin and Franklin counties identified by location number.

**Figure S2.** Relative abundance of top 10 identifiable phyla shown across all samples. Location names (Location 1 through Location 6) are abbreviated as L1 through L6.

**Figure S3.** Relative abundance of top 10 genera (identifiable at least to family level) shown across all samples. Location names (Location 1 through Location 6) are abbreviated as L1 through L6.

**Figure S4**. Principal Coordinates Analysis of all samples based on Bray-Curtis distance. Location names (Location 1 through Location 6) are abbreviated as L1 through L6. Substrate samples are clearly separated from the tardigrade samples along Axis 1. On Axis 2, samples from 2020 are generally clustered away from 2019 samples. Samples from different locations and from lichen and moss overlap.

## Supplemental Table Legends

**Table S1.** Mothur classification and decontam score for all OTUs with more than 10 reads in experimental samples. OTUs with decontam score below the 0.25 threshold are marked as contaminants.

**Table S2.** BLAST results for OTUs with more than 10 total reads in experimental samples. BLAST+ v2.11.0 was used to query representative sequences for each OTU against a database generated from the NCBI 16S RefSeq collection. The 15 hits with the lowest E-values are given for each OTU.

**Table S3.** RDP Classifier results for OTUs with more than 10 total reads in experimental samples. Representative sequences for each OTU were uploaded to the RDP Classifier webtool (https://rdp.cme.msu.edu/classifier/classifier.jsp), version 2.11 using 16S rRNA training set 18.

**Table S4.** Relative abundance of all analyzed OTUs in each sample.

**Table S5.** Relative abundance of each OTU by level of Contrast 1 (Location).

**Table S6**. Relative abundance of each OTUs in moss and in lichen (*i.e*., at each level of Contrast 2).

**Table S7**. Relative abundance of each OTU in tardigrades and in their lichen substrate (*i.e*., at each level of Contrast 3).

**Table S8**. Relative abundance of each OTU in tardigrades in 2019 and in 2020 (*i.e*., at each level of Contrast 4).

**Table S9.** Differentially abundant and variable taxa across contrasts. Corncob was used to identify significantly (Benjamini-Hochberg corrected *P* value < 0.05) differentially abundant and variable taxa across four contrasts of interest. Genus names are presented including their phylum, class, order, and family names to prevent ambiguities.

**Table S10.** Relative abundance of top 10 identifiable phyla across levels of each contrast.

**Table S11.** Relative abundance of top 10 genera (identifiable at least to family level) across levels of each contrast.

