## Supplementary figures and images for "Tardigrade community microbiomes in North American orchards include putative endosymbionts and plant pathogens"

### Figure S1

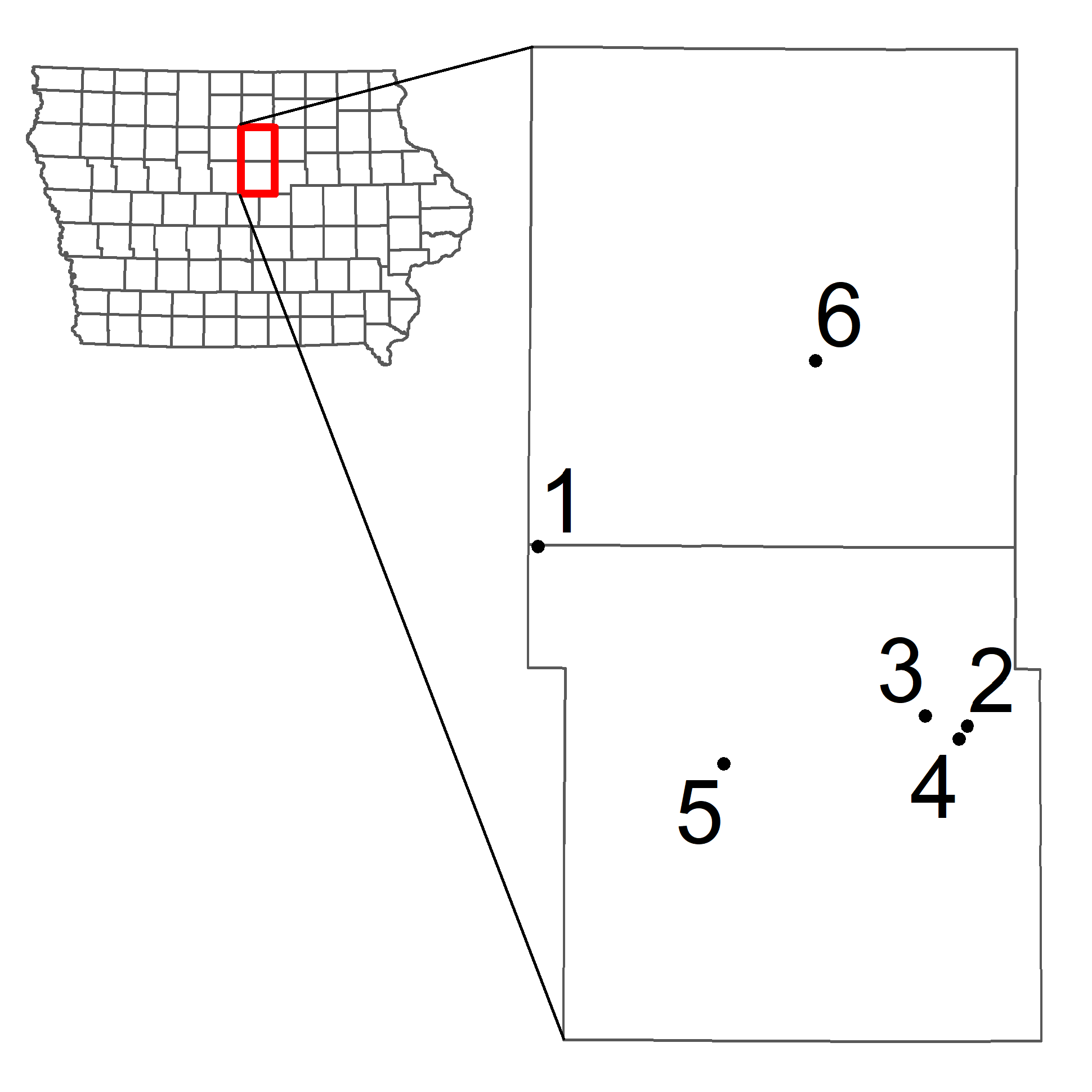

### Figure S2

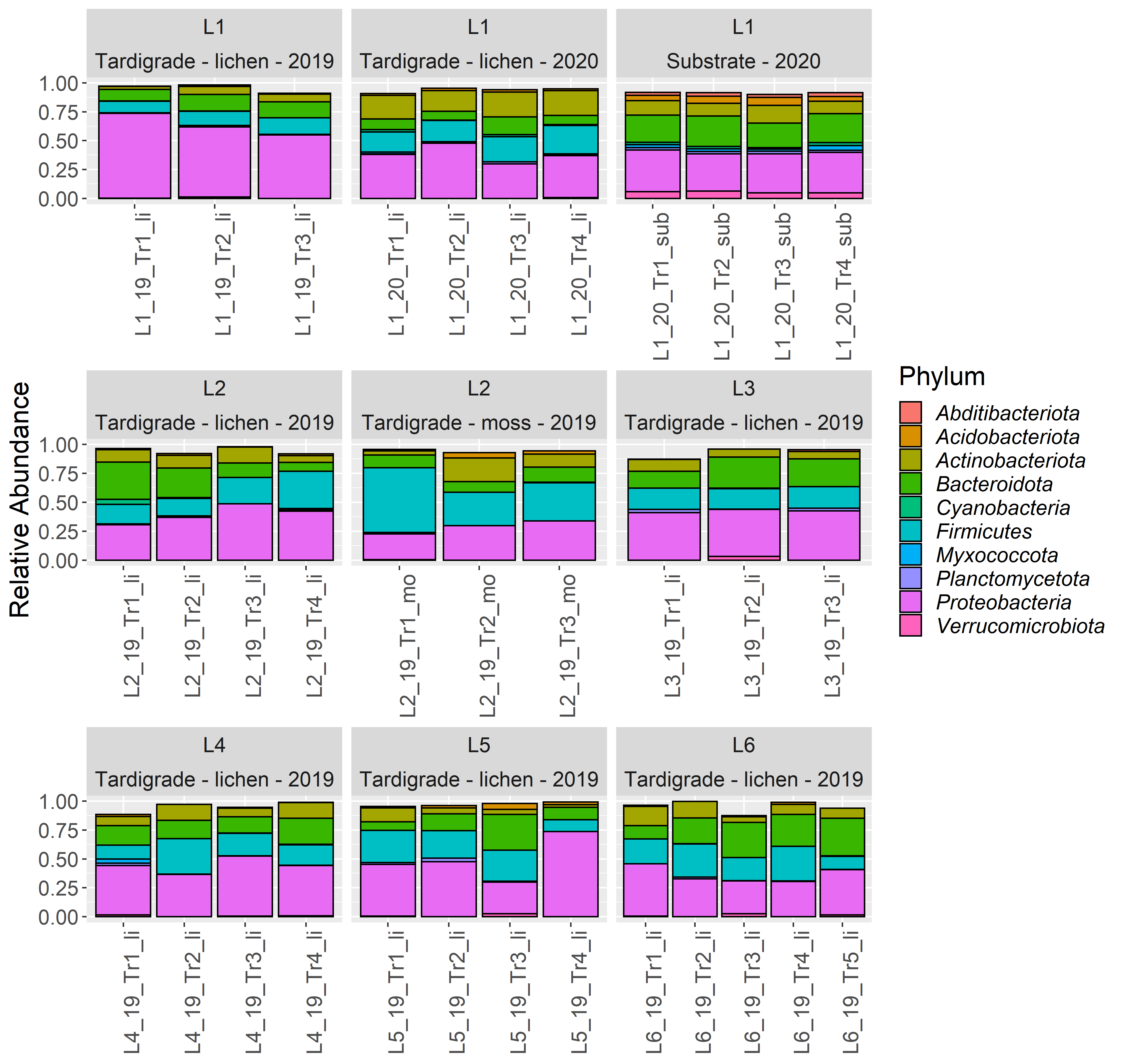

### Figure S3

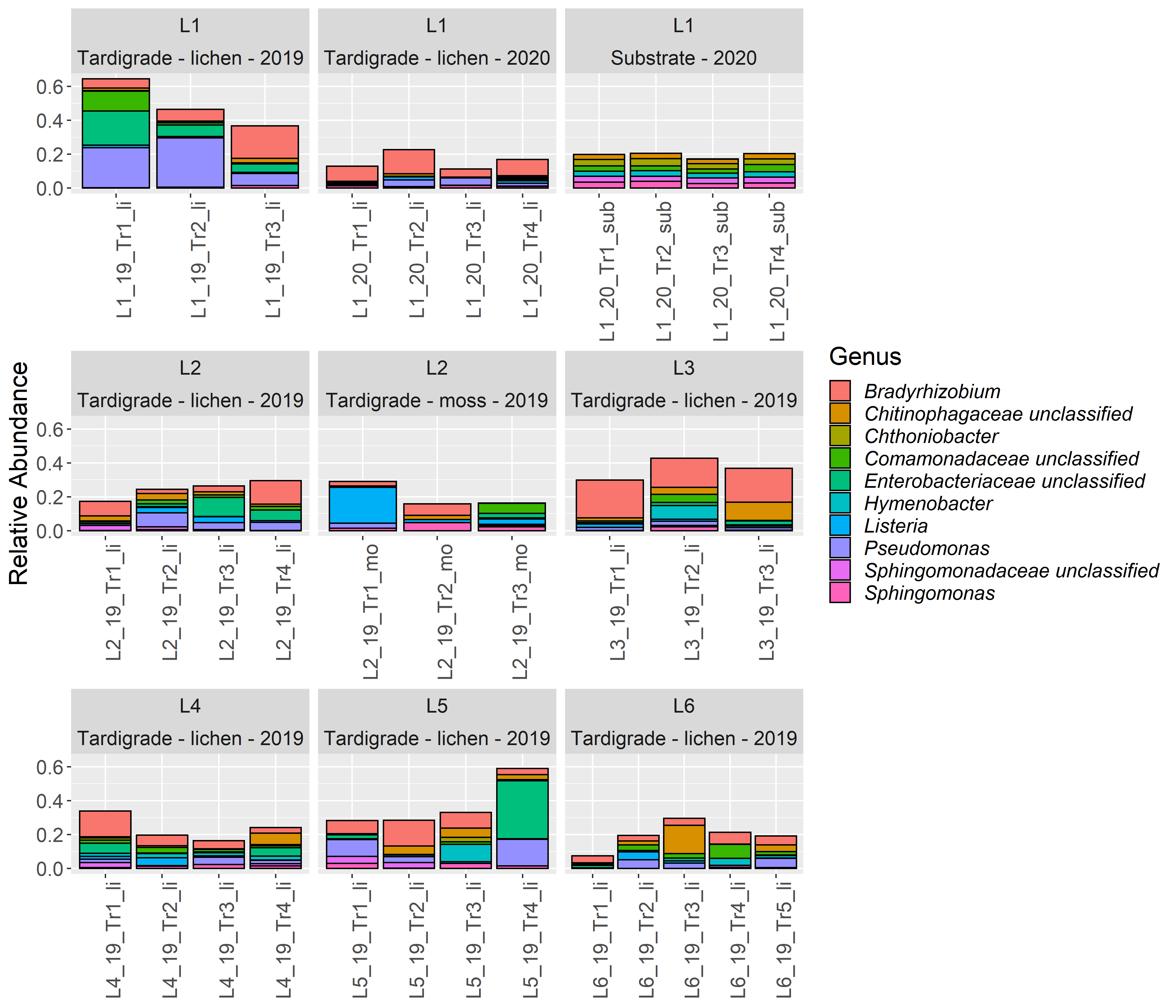

### Figure S4

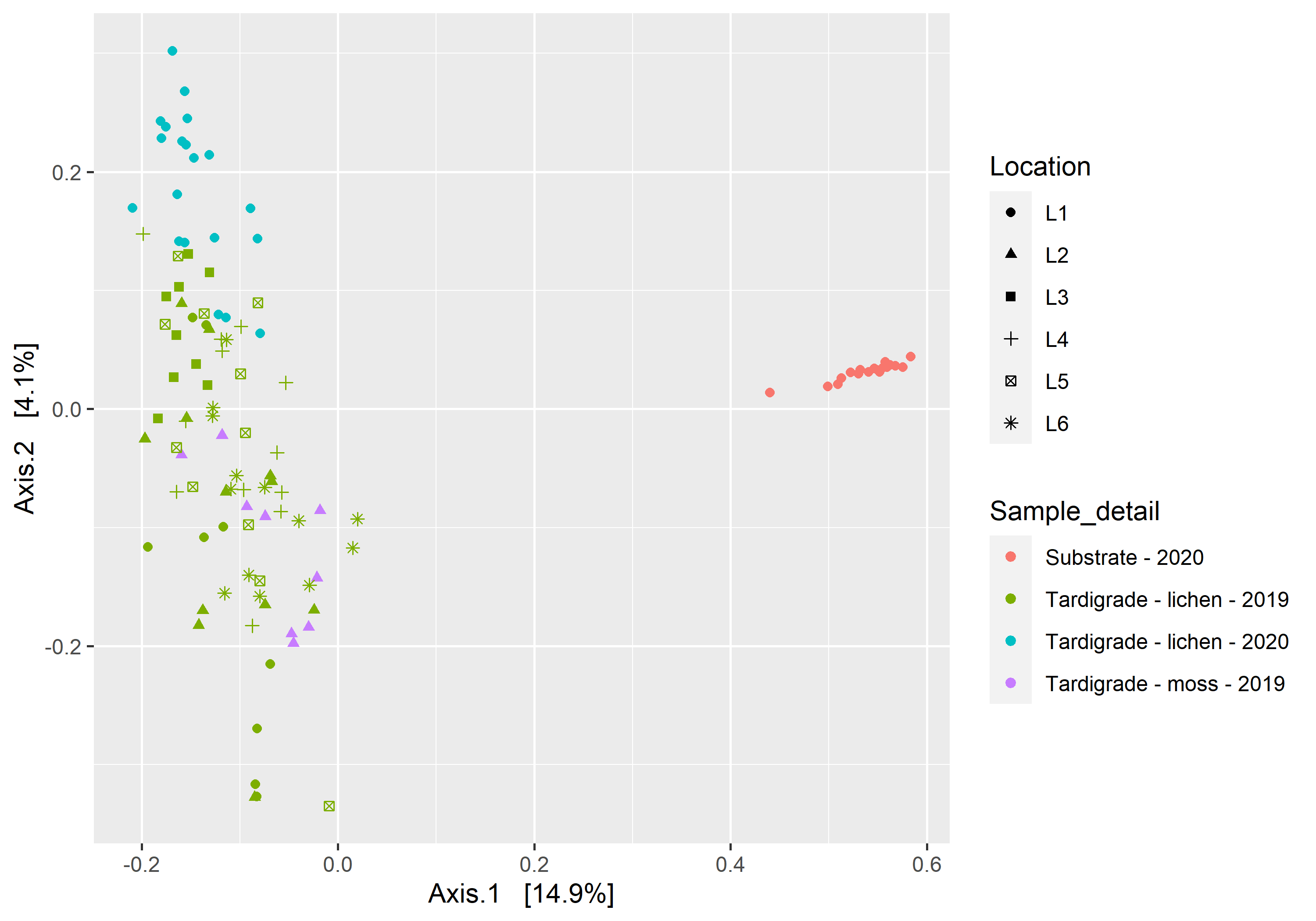
